# Self-amplifying mRNA expression is governed by mitochondrial machinery and ACSL4

**DOI:** 10.1101/2025.06.13.659366

**Authors:** Lara F. Tshering, Chong Wang, Qing Xiang, Sharmistha Kundu, Alicia Solórzano, Javier Chaparro-Riggers, Laura Lin, Azadeh Paksa

## Abstract

RNA-based vaccines offer a superior efficacious and economic approach over traditional vaccines. However, current dose requirement and short half-life of conventional modified mRNA (modRNA) may hinder the development of more effective vaccines. Self-amplifying mRNA (saRNA) is a modality that has the potential to address these limitations by reducing delivery dosage and enhancing the duration of expression over modRNA. Despite marked success in preclinical studies, saRNA vaccines have thus far underperformed in most clinical trials. We hypothesized that non-optimal human cellular context limits saRNA expression, and that elucidating the factors underlying saRNA expression would be key in developing saRNA as a viable therapeutic modality. To identify factors involved in the regulation of saRNA, we performed a quasi-genome-wide CRISPR knockout screen, which revealed that saRNA expression in human cells is linked to mitochondrial function and governed by ACSL4. We validated these findings through pharmacological intervention and single gene editing, demonstrating that ACSL4-mediated regulation is unique to saRNA and does not impact modRNA. Moreover, we show that modulating ACSL4 leads to improved saRNA expression across multiple cell types. We demonstrate for the first time that mitochondrial function and ACSL4 play key roles in saRNA expression, providing insight for the development and implementation of saRNA as a therapeutic modality.

## INTRODUCTION

As an essential public health tool, optimizing the development and distribution of vaccines is a top priority globally. RNA-based vaccines offer a superior approach over traditional modalities through therapeutic efficacy, rapid development timeline, highly scalable manufacturing, and well-tolerated safety profiles^1–3^. The power of non-replicating conventional modified mRNA (modRNA) vaccines is clearly demonstrated by their unprecedented success in preventing severe disease and minimizing infections by severe acute respiratory syndrome coronavirus 2 (SARS-CoV-2)^4,5^. Comirnaty (Pfizer, BioNTech) and Spikevax (Moderna) played an instrumental role in combatting the COVID-19 pandemic, opening the door for developing similar modRNA vaccines for other infectious diseases and therapeutics. Nonetheless, challenges currently faced by conventional modRNA vaccines include dose requirement and short half-life of the vaccine, which limit the utilization of this modality for therapeutics^1,2^. As an alternative RNA-based modality, self-amplifying mRNA could potentially address these limitations.

Self-amplifying mRNA (saRNA) is derived from the genome of alphaviruses such as the Venezuelan equine encephalitis (VEEV) and Sindbis viruses (SINV)^1,2^. Comprised of the same basic structure as conventional mRNA vectors, saRNA are distinguished by the presence of a subgenomic promoter and a large open-reading frame encoding four nonstructural proteins^2,3^. The gene of interest (GOI) replaces the viral genome, rendering saRNAs incapable of producing viral structural proteins and therefore live virus^3^. Translation of the nonstructural proteins give rise to the functional component of the viral replicon, an RNA-dependent RNA polymerase (RDRP) complex, upon engaging of a host cell’s ribosome. The RDRP complex transcribes more copies of genomic RNA and amplifies subgenomic mRNA to drive expression of the GOI. This results in high levels of sustained GOI expression, offering a potent modality for vaccines and therapeutics by minimizing the usage of delivery materials while achieving more durable expression and protection with fewer injection regimens compared to modRNA^2,3,6,7^.

Preclinical *in vitro* and *in vivo* studies have demonstrated that saRNA vaccines outperform modRNA and traditional DNA or protein modalities in multiple disease types^1,6,8–10^. However, to date the majority of efforts to develop saRNA as a therapeutic and vaccine modality have turned underwhelming results, with few exceptions^1,7,11–14^. The lack of translation from preclinical to clinical success could be due to a lack of host cellular context of the factors that control saRNA expression, specifically within human immune cells. *In vitro* and *in vivo* studies have documented a strong interferon response to saRNA^15,16^, and while there is evidence that these same interactions exist in humans, they are currently poorly understood^3^. We hypothesized that bridging this gap in knowledge and elucidating the cellular factors underlying saRNA expression would be key in developing saRNA as a viable therapeutic modality.

To identify these factors, we used pooled Clustered Regularly Interspaced Short Palindromic Repeats knockout (CRISPR-KO) screening to study phenotypes at a quasi-genomic scale^17,18^. We identified mitochondrial function and ferroptosis as key biological processes involved in the expression of saRNA in multiple human cell types, including immune cells, with Acyl-CoA Synthetase Long Chain Family Member 4 (ACSL4) emerging as a key factor in controlling saRNA expression. Furthermore, we show that these factors specifically regulate saRNA, with no effect on modRNA expression, rationalizing the discrepancy between the success of modRNA and saRNA in clinical trials. Additionally, we show that increasing ACSL4 in multiple cell types improves saRNA expression, highlighting one potential strategy to unleash the power of saRNA as a therapeutic modality. To our knowledge, this study represents the first evidence of a link between ACSL4 and saRNA in human cells.

## RESULTS

### Quasi-genome wide CRISPR screen reveals that saRNA expression is linked to mitochondrial function and ferroptosis

To identify the underlying factors involved in saRNA expression in human cells, we pursued pooled CRISPR-KO screening, leveraging a previously established genome-wide pooled guide RNA (gRNA) library^19^ (Figure 1A). We first produced non-modified saRNA comprising standard eGFP and HiBit^20,21^ reporters (saGFP). We co-transfected our saGFP with a 5-Methoxyuridine-5’-Triphosphate (5-Methoxy-UTP) modified mRNA carrying mCherry reporter (mod-mCherry) into standard 293T cells at various ratios in order to establish a cellular model paradigm and transfection system for the screen challenge^17,18^ (Figure 1B, Supplemental Figure 1A). Co-transfection was devised to single out hits unique to saRNA expression while excluding hits associated with general RNA transfection, thereby identifying key factors needed to adapt saRNA for clinical use. Upon evaluating the proportion of dual versus single fluorophore expressors, we observed that dual-positive populations varied based on RNA concentration ratios but no singly GFP+ population existed in any co-transfection condition; all GFP+ cells were dual expressors (Figure 1B). This observation suggests that while modRNA is consistently and reliably expressed in 293T cells, even among wild type cells not all are able to express saRNA. Based on these data, in all following experiments we used a 1:0.5 ratio of modRNA to saRNA (half “dose” of saRNA), as it showed the most equitable distribution between mCherry+ and dual-positive populations.

**FIGURE 1:**
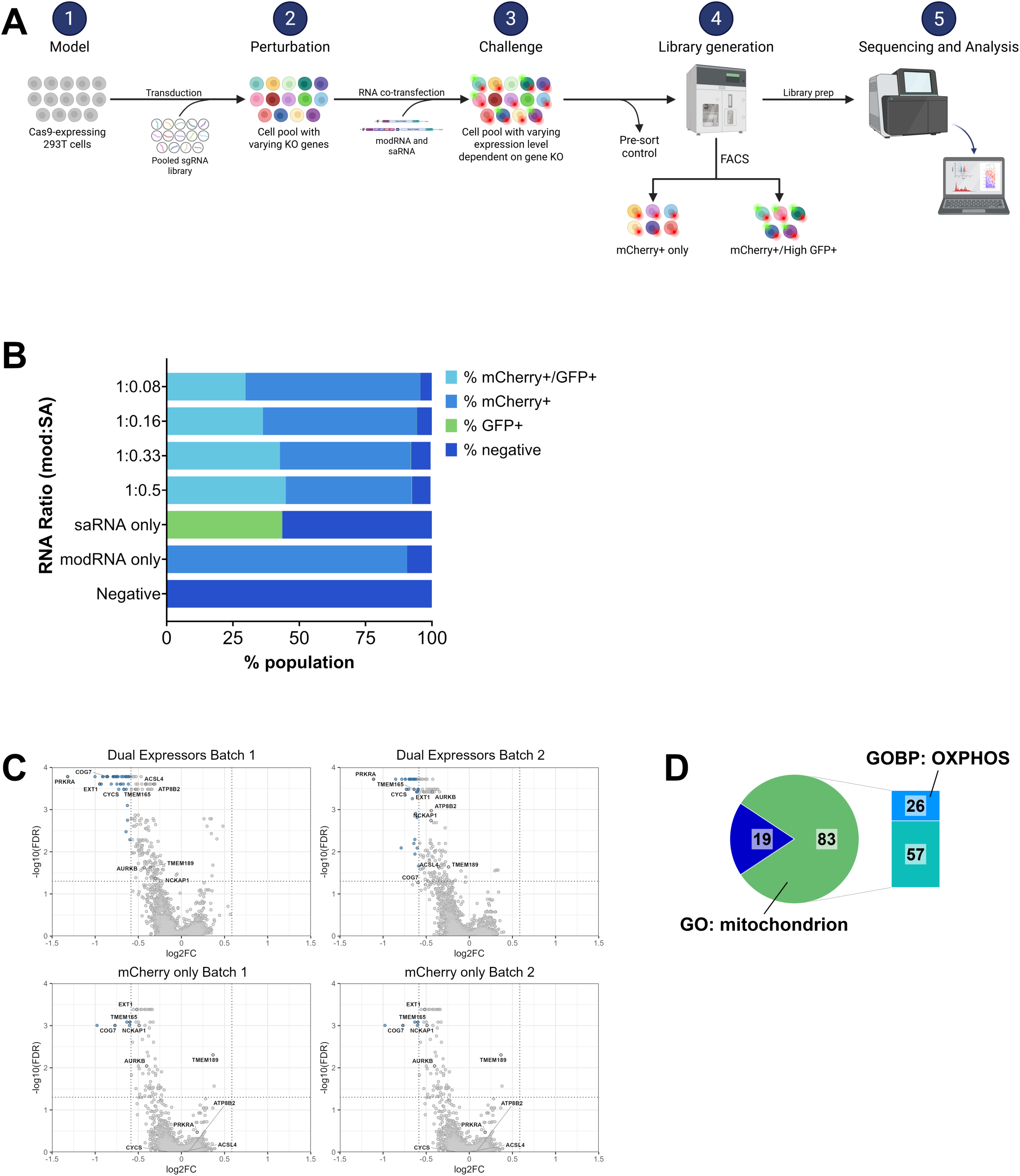
Quasi-genome wide CRISPR screen identifies mitochondrial function as a key biological process in saRNA expression. A) Schematic for CRISPR-KO screen detailing the plan for each typical step of a CRISPR screen including model, perturbation, screen challenge, library generation, and analysis. Created in BioRender (biorender.com/u36hl45). B) 293T cells overnight co-transfected with saGFP and mod-mCherry at indicated ratios in 96 well format. mod-mCherry concentration was kept constant at 50 ng/well while saGFP varied from 4 to 25 ng/well. C) Volcano plots for each screen sample displaying log_2_(fold change) as calculated by MAGeCK. In the top left quadrant for each plot, genes highlighted in blue exhibit −log_10_(false discovery rate) > 1 and log_2_(fold change) < −0.5. Five named genes highlighted are common across all samples, appearing significant on all plots; the other five named genes highlighted are unique to Dual Expressors, only appearing significant in these samples. Essential genes integrated into each pool as an internal control performed as expected, indicating successful Cas9 editing efficiency (data not shown). D) Hits unique to Dual Expressor samples and suspected necessary factors for saRNA expression are primarily associated with mitochondrial function. General mitochondria (GO: mitochondrion) and oxidative phosphorylation (GOBP: OXPHOS) gene lists are taken from Gene Ontology and Gene Ontology Biological Process databases.

To evaluate whether this effect was unique to 293T cells or consistent across human cell types, we co-transfected saGFP and mod-mCherry into Jurkat cells, fibroblasts, and primary normal human dendritic cells (NHDCs). Jurkat cells were used as they have previously shown robust expression of saRNA^22^. Fibroblasts were used as a non-immune cell with an intact interferon response that was previously shown to express saRNA^23^. NHDCs were used as a proxy for *in vivo* target cells due to their key role in developing adaptive immune responses^24^. While fluorophore distribution between populations varied based on cell type, we confirmed the lack of singly GFP+ population across all cell types (Supplemental Figure 1B). Taken together, these observations suggest that saRNA expression, and likely the factors involved in its regulation, is highly selective and variable in different cell types. Due to the inconsistency of dual positivity rate in the additional cell types tested (data not shown), we opted to pursue our screen in 293T cells, as an immune-incompetent line, which would allow an understanding of fundamental machineries underlying saRNA expression separately from immune processes.

To pursue our quasi-genome-wide screen, we used three quarters of a previously established gRNA library^19^. We generated stable Cas9-expressing 293T (293T-Cas9) cells with sufficient Cas9 editing efficiency (Supplemental Figure 1C). gRNA libraries were introduced to 293T-Cas9 cells via lentivirus for the perturbation step of the screen (Figure 1A). For the screen challenge, pooled cell libraries were co-transfected with saGFP and mod-mCherry, followed by FACS, library prep, and sequencing (Figure 1A). Two separate populations were collected via FACS (Supplemental Figure 1D): 1) high GFP expressing dual positive cells, representing factors associated with saRNA expression; 2) single mCherry expressors, representing factors associated with modRNA expression. To minimize false-positive hits and increase robustness in hit identification and library coverage, screen challenge was performed in duplicate on consecutive days (Figure 1C, Supplemental Figure 1D). Pre-sort samples within each respective replicate batch were collected for comparison.

To identify necessary factors for saRNA expression, we focused on “negative” hits: gRNAs identified as depleted in each sample and therefore suspected necessary factors of expression. Only gRNAs depleted in both replicates were selected as “true” hits, resulting in three subsets: 1) hits common across all samples; 2) dual mCherry+/GFP+ specific hits; 3) and single-mCherry+ specific hits (Supplemental Table 1). The latter, made up of hits unique to modRNA expression, was the smallest subset (n=7) and consisted of random, unrelated genes; this list was disregarded. We hypothesized that hits common across all samples (n=11) were implicated in general lipofectamine transfection. Indeed, all identified genes were associated with heparan sulfate, an essential factor for lipofectamine transfection in CHO cells^25^. These data suggest our internal controls and screen design are sufficient to produce relevant hits. Our main subset of interest, dual mCherry+/GFP+ specific hits, represent suspected necessary factors for saRNA expression (Table 1). This was the largest subset (n=102) and mainly consisted of genes associated with mitochondrial function (Figure 1D, Table 1). Acyl-CoA Synthetase Long Chain Family Member 4 (ACSL4) was among the top ranked hits. ACSL4 stood out to us as both a necessary factor of ferroptosis^26^ (an iron-dependent form of cell death^27^) and single-stranded RNA virus infection^28^, highlighting a potential crossover between saRNA expression and ferroptosis. Furthermore, mitochondrial function and ferroptosis are well-established to be inextricably linked^26,29–34^, while ferroptotic pathways are functional only in a subset of immune cells^32^. Together, this suggests that many factors involved in saRNA expression in human cells are associated with mitochondrial function and ferroptosis, presenting a rational hypothesis for the difficulty of saRNA expression in certain human cell types.

### Pharmacological intervention and single gene editing of top hits reveal dependency on mitochondrial function and ferroptosis is unique to saRNA

To evaluate how mitochondrial function and ferroptosis influence saRNA expression, we examined the functional effect of these biological processes on transfection *in vitro*. We pre-treated 293T cells with electron transport chain (ETC) inhibitors rotenone (Complex I) or antimycin A (Complex III)^31^, followed by transfection with either modRNA or saRNA carrying identical GFP and HiBit^20,21^ reporters (modGFP or saGFP). For both treatments, saRNA expression was significantly impaired when compared to its corresponding negative control across all concentrations (Figure 2A, Supplemental Figure 2A). Antimycin A had no significant effect on modRNA expression (Figure 2A-B). Although rotenone reduced modRNA expression across all concentrations, the loss of signal was significantly more pronounced in saRNA transfected cells (Supplemental Figure 2A-B), suggesting that saRNA is more sensitive to disruption in mitochondrial activity. Next, we demonstrated that pre-treatment of 293T cells with ferrostatin-1 (Fer-1), an antioxidant and ferroptosis inhibitor^27^, significantly decreased saRNA expression at high concentrations while having no effect on modRNA (Figure 2C). Together, these findings validate the impact of mitochondrial function and ferroptosis as key biological processes in saRNA expression in human cells.

**FIGURE 2:**
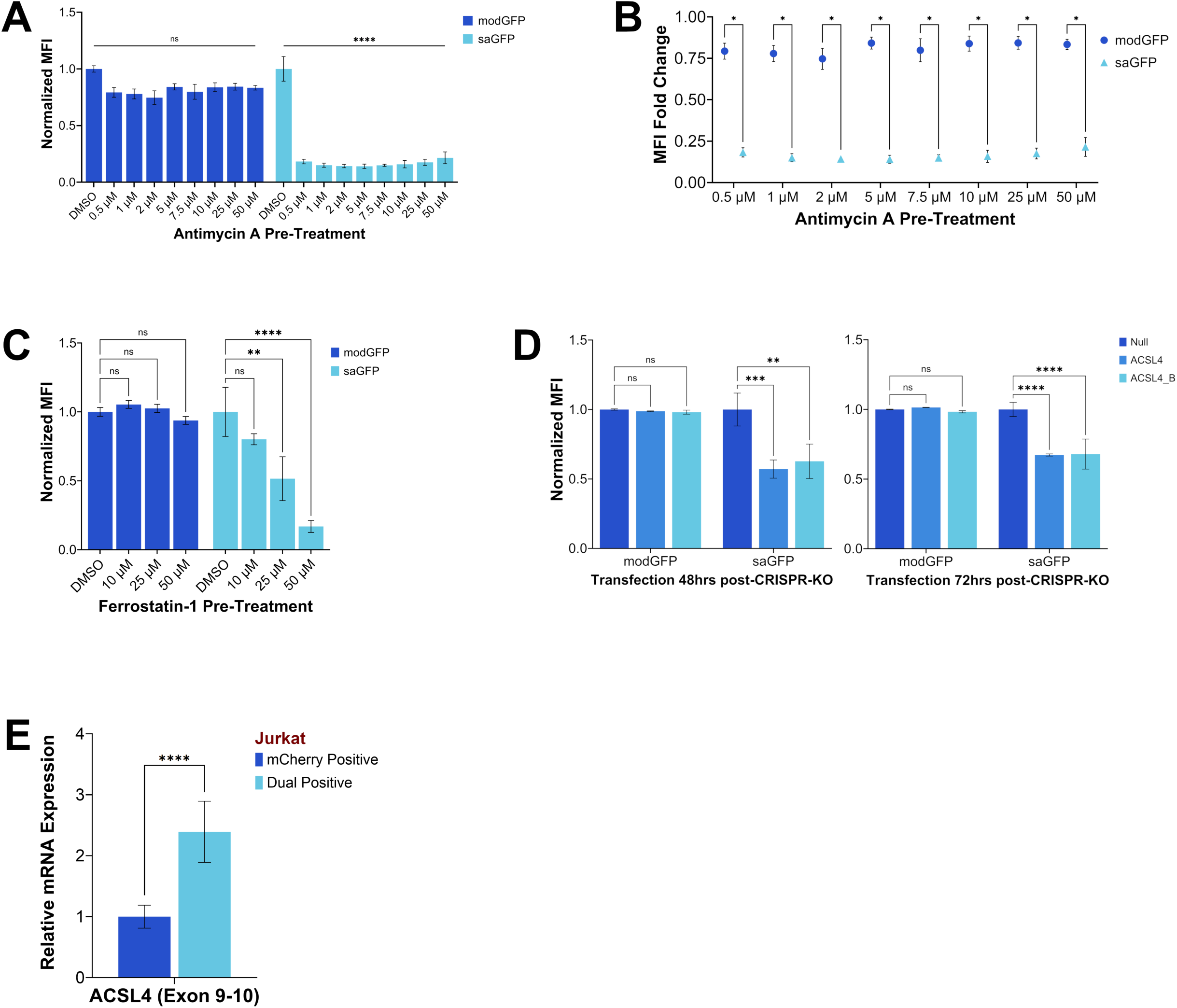
Pharmacological intervention and single gene editing demonstrate dependency on mitochondrial function and ferroptosis is unique to saRNA expression. A) 293T cells pre-treated with antimycin A overnight at the concentrations indicated before transfection with modGFP or saGFP. Each well received a standard dose of RNA (120 ng/well in 96 well format). MFI was read by HiBit assay after four hours and normalized to the negative control of each modality (DMSO). n=3 biological replicates. B) Comparison of fold change in MFI for each concentration of antimycin A pre-treatment. saRNA MFI is significantly more impacted at all concentrations. C) 293T cells pre-treated with ferrostatin-1 (Fer-1) overnight at indicated concentrations before transfection with modGFP or saGFP show Fer-1 only abrogates saRNA expression. Each well received a standard dose of RNA (120 ng/well in 96 well format). MFI was read by HiBit assay after four hours and normalized to the negative control of each modality (untreated/DMSO). n=3 biological replicates. D) 293T-Cas9 cells transfected with two unique ACSL4-targeting guide RNAs show ACSL4 dependency is unique to saRNA expression. KO cells were subcultured for 48 (left) or 72 (right) hours prior to transfection with modGFP or saGFP. Each well received a standard dose of RNA (120 ng/well in 96 well format). MFI was read by HiBit assay after four hours and normalized to the negative control of each modality (gRNA null/normal 293T-Cas9 cells). n=3 biological replicates. E) Jurkat cells co-transfected with saGFP and mod-mCherry overnight show dual expressors naturally exhibit higher levels of ACSL4 expression as measured by qPCR. Jurkat cells were nucleofected using Lonza 4D-Nucleofector™ 96-well Unit with a cell density of 5e5 cells per well; co-transfection wells received 1000 ng modRNA and 500 ng saRNA per well. Single modality and H_2_O nucleofection conditions were used as gating controls for FACS. 2^-ΔΔ*CT*^ was calculated using mCherry single expressors as the control.

To validate the top candidates of key genes required for saRNA expression, we generated single-gene CRISPR-KO cells using our 293T-Cas9 cells. Three top genes from our saRNA specific list were selected; pre-designed gRNAs targeting these genes (Synthego, IDT) were used (Supplemental Table 2). 293T-Cas9 cells were transfected with each gRNA to generate four KO lines. Each line was expanded for 48 or 72 hours prior to modRNA or saRNA delivery. Although no effect was observed on modRNA expression in all KO lines, saRNA expression was significantly impeded regardless of time point (Figure 2D, Supplemental Figure 2C) despite a suboptimal editing efficiency (≤70%) in all lines (data not shown). This supports our screen’s successful identification of factors unique to saRNA. Furthermore, disruption of our top candidate ACSL4 with multiple independent gRNAs showed a significant decrease in saRNA expression with no observable impact on modRNA (Figure 2D). qPCR analysis in Jurkat cells co-transfected with saGFP and mod-mCherry revealed significantly increased expression of ACSL4 in dual-expressing cells over single expressors (Figure 2E). These data confirm that ACSL4 is a key factor unique to saRNA expression in human cells, regardless of immune competency, and most likely acts through its role in mitochondrial function and ferroptosis^26,35^.

### Early events on the mitochondrial-ferroptotic axis are involved in saRNA expression

We next sought to identify the processes on the mitochondrial-ferroptotic axis that govern saRNA expression. 1S,3R-RSL3 (RSL3) induces ferroptosis through inhibition of GPX4^27,29,31^, acting downstream of mitochondrial involvement in this axis (Figure 3A). While mitochondria are essential in cysteine-deprivation-induced ferroptosis, their involvement is dispensable for GPX4-related ferroptosis^31^. As inhibition of ferroptotic signaling diminished saRNA expression, we hypothesized that if ferroptosis itself was the key process in saRNA machineries, inducing ferroptosis would have the opposite effect. To test this, we pre-treated 293T cells with RSL3 prior to RNA transfection, inhibiting GPX4 and inducing ferroptosis. At high concentrations, RSL3 pre-treatment significantly diminished RNA expression regardless of modality, and saRNA was impacted at all concentrations (Figure 3B), likely due to increased cytotoxicity. Our data therefore suggest that early events on the mitochondrial-ferroptotic axis are the critical processes in saRNA expression (Figure 3A, highlighted in blue). To confirm early events govern saRNA expression, we investigated the effect of transfection on lipid peroxidation in 293T cells. As lipid peroxidation occurs as a result of byproducts of normal mitochondrial function^31^, we rationalized that if saRNA machineries were involved in this phase of the mitochondrial-ferroptotic axis, successful saRNA transfection would result in higher rates of lipid peroxidation. To detect lipid peroxidation, we stained cells with BODIPY lipid peroxidation sensor (Invitrogen) after modRNA or saRNA transfection. Across all concentrations, there was no significant impact on detected oxidation regardless of RNA modality (Figure 3C). We therefore conclude that saRNA is intrinsically linked to mitochondrial activity and upstream events on the mitochondrial-ferroptotic axis (Figure 3A).

**FIGURE 3:**
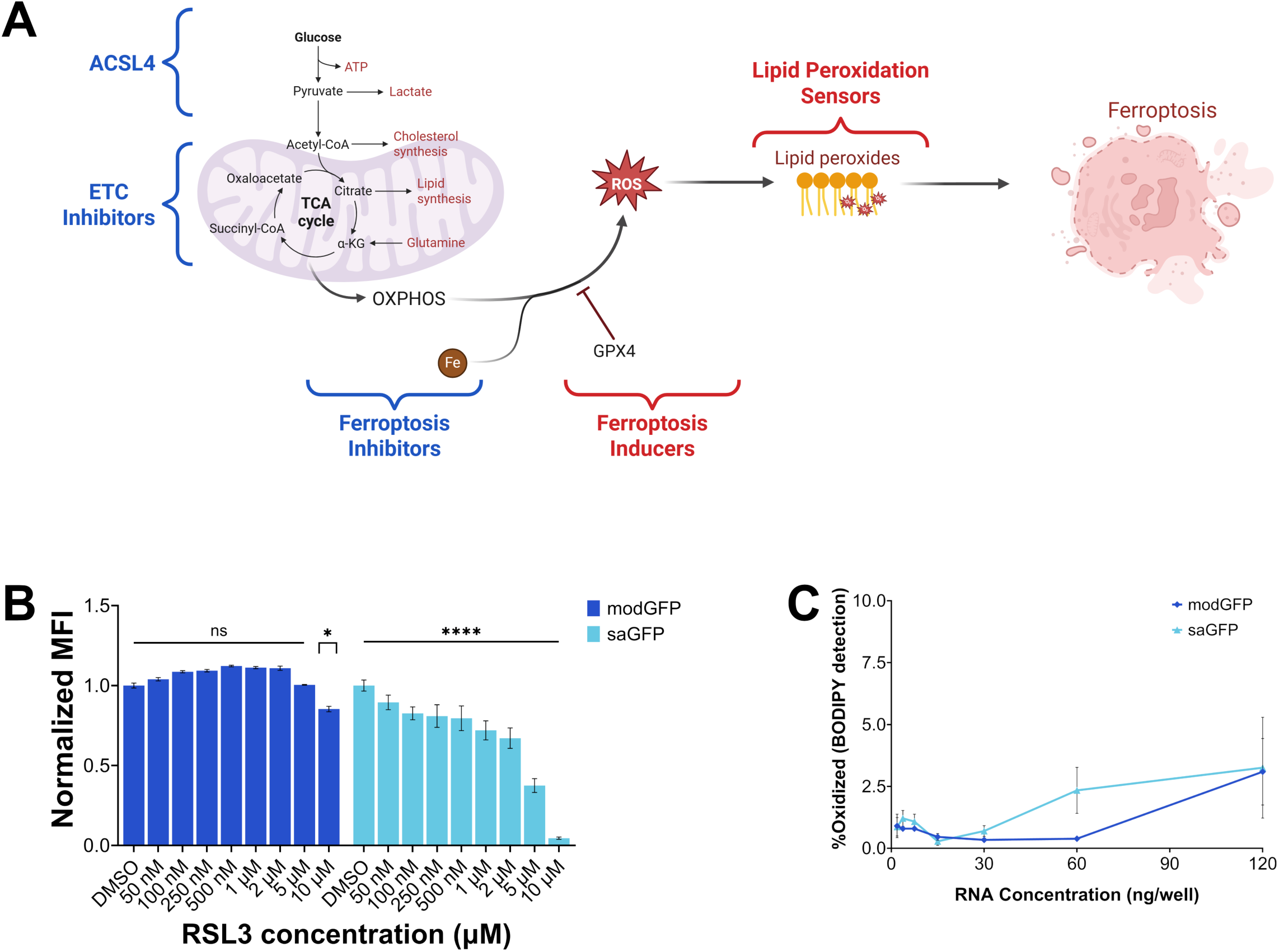
Early events on the mitochondrial-ferroptotic axis are involved in saRNA expression. A) Schematic representing the mitochondrial-ferroptotic axis and the interaction between these biological processes. ACSL4 activity, electron transport chain (ETC) inhibitors, and ferroptosis inhibitors that have demonstrated a clear impact on saRNA expression are highlighted in blue, demonstrating their location upstream in this axis. Ferroptosis inducers and lipid peroxidation are highlighted in red and located at downstream points in this axis. Created in BioRender (biorender.com/5rmsile). B) 293T cells pre-treated with RSL3 for one hour at indicated concentrations show RNA expression is sensitive to high concentrations of RSL3 regardless of RNA modality. Each well received a standard dose of modGFP or saGFP (120 ng/well in 96 well format). MFI was read by HiBit assay after four hours and normalized to the negative control of each modality (untreated/DMSO). n=3 biological replicates. C) Lipid peroxidation detected by BODIPY dye oxidization shows no effect on lipid peroxidation post-transfection regardless of RNA modality. Each well received a standard dose of modGFP or saGFP (120 ng/well in 96 well format) for overnight transfection. Cells were analyzed by flow cytometry; debris, dead cells, doublets, and GFP− cells were gated out by FlowJo to observe oxidation in GFP+ cells only. n=3 biological replicates.

### ACSL4 activation leads to improved saRNA expression *in vitro*

Focusing on early events, we turned our attention to ACSL4, our top gene of interest and a factor acting upstream of mitochondrial activity (Figure 3A). We first sought to activate ACSL4 expression by CRISPRa, incorporating mCherry-dCas9^36^ into 293T cells to do so. Following selection by FACS, cells were transfected with ACSL4-targeting gRNAs and incubated for 48 hours prior to modRNA and saRNA transfection. Despite only a modest but significant increase in ACSL4 expression in ACSL4-activated cells (Figure 4A), saRNA expression increased almost 3-fold while modRNA expression remained unchanged (Figure 4B). To control for potential off-target CRISPR effects, we generated 293T cells overexpressing ACSL4 using pre-designed ORF vectors (OriGene) delivered via lentivirus. This led to a marked increase in ACSL4 (Supplemental Figure 3A); we observed no changes in cell viability, suggesting that increased ACSL4 levels do not induce cytotoxicity (Supplemental Figure 3B). Overexpression of ACSL4 resulted in improved saRNA expression as shown by significant increase in the percentage of GFP+ cells transfected with saGFP (Supplemental Figure 3C), in addition to the percentage of dual expressors in our co-transfection assay (Supplemental Figure 3D). We therefore concluded that activation of ACSL4 leads to increased saRNA expression *in vitro*.

**FIGURE 4:**
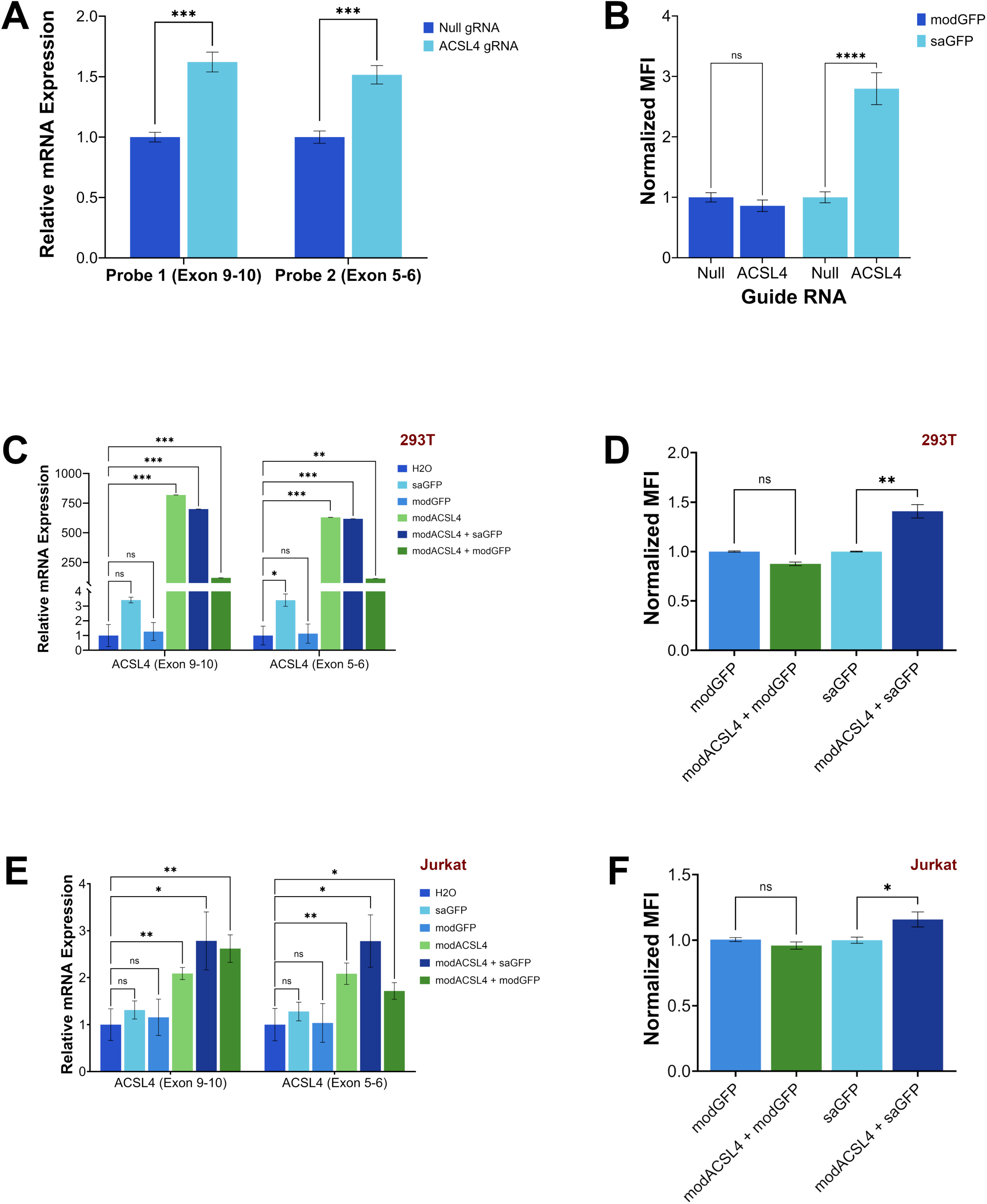
ACSL4 activation *in vitro* improves saRNA expression but not modRNA. A) ACSL4 expression as read by qPCR with two unique probes targeting different exons shows successful activation of ACSL4 via CRISPRa. Gene expression assay was conducted concurrently with RNA IVE. 2^-ΔΔ*CT*^ was calculated using normal 293T-dCas9 cells (no gRNA/normal 293T-dCas9 cells) as the control. B) ACSL4-activated cells show increased saRNA expression. CRISPRa cells were subcultured for 48 hours after gRNA transfection before being transfected with modGFP or saGFP. Each well received a standard dose of RNA (120 ng/well in 96 well format). MFI was read by HiBit assay after four hours and normalized to negative control of each modality (gRNA null/normal 293T-dCas9 cells). n=3 biological replicates. C) Transfection with modACSL4 successfully increases ACSL4 expression in 293T cells as read by qPCR. 293T cells were transfected with either modGFP, saGFP, modASCL4, or co-delivered modACSL4 with a GFP carrying RNA overnight in a 96 well format. Each single modality transfection received 100 ng/well RNA. Co-transfection with saRNA received 50 ng modRNA/well combined with 25 ng saRNA/well. Co-transfection of two modRNAs received 50 ng of each modRNA per well. All cells that received modACSL4 (either alone or co-delivered) exhibited significantly higher ACSL4 expression with both unique probes. 2^-ΔΔ*CT*^ was calculated using non-transfected 293T cells (H_2_O negative control) as the control. D) MFI of 293T cells with co-delivered modACSL4 and GFP carrying RNA show increased saRNA expression with no effect on modRNA expression. 293T cells were transfected with either modGFP, saGFP, modASCL4, or co-delivered modACSL4 with a GFP carrying RNA overnight in a 384 well format. Each single modality transfection received 50 ng/well RNA. Co-transfection with saRNA received 25 ng modRNA/well combined with 12.5 ng saRNA/well. Co-transfection of two modRNAs received 25 ng of each modRNA per well. MFI of each co-transfection was normalized to its single modality counterpart. n=4 biological replicates. E) Transfection with modACSL4 successfully increases ACSL4 expression in Jurkat cells as read by qPCR. Jurkat cells were nucleofected with either modGFP, saGFP, modASCL4, or co-delivered modACSL4 with a GFP carrying RNA overnight using Lonza 4D-Nucleofector™ 96-well Unit with a cell density of 5e5 cells per well. Each single modality transfection received 1000 ng/well RNA. Co-transfection with saRNA received 500 ng modRNA/well combined with 250 ng saRNA/well. Co-transfection of two modRNAs received 500 ng of each modRNA per well. All cells that received modACSL4 (either alone or co-delivered) exhibited significantly higher ACSL4 expression with both unique probes. 2^-ΔΔ*CT*^ was calculated using non-transfected Jurkat cells (H_2_O negative control) as the control. F) MFI of Jurkat cells with co-delivered modACSL4 and GFP RNA show increased saRNA expression with no effect on modRNA expression. Jurkat cells were nucleofected as described in 4E. Cells were analyzed by flow cytometry; debris, dead cells, doublets, and GFP− cells were gated out by FlowJo to observe MFI in GFP+ cells only. MFI of each co-transfection was normalized to its single modality counterpart. n=3 biological replicates.

To replicate this effect in a more clinically relevant manner, we devised a co-transfection approach with our saGFP and an ACSL4-targeting modRNA (modACSL4) acting as a therapeutic adjuvant. Starting with 293T cells, we observed significant increase of ACSL4 expression in all samples that received modACSL4 (Figure 4C) with no major impact on cytotoxicity (Supplemental Figure 4A). As anticipated, co-transfection with modACSL4 improved saRNA expression significantly (Figure 4D, Supplemental Figure 4B), but had no effect on modRNA (Figure 4D). We next repeated this co-transfection in Jurkat cells, fibroblasts, and NHDCs. modACSL4 consistently increased ACSL4 expression across all cell types (Figure 4E, Supplemental Figure 4C, Supplemental Figure 4E). Enhanced ACSL4 levels were sufficient to modestly but significantly increase saRNA expression in all cell types while having no effect on modRNA (Figure 4F, Supplemental Figure 4D, Supplemental Figure 4F). These data demonstrate ACSL4 activation as a safe and effective adjuvant for saRNA, presenting a strong rationale for *in vivo* studies of co-delivering modACSL4 and saRNA as a therapeutic modality.

## DISCUSSION

To date, the limited success of saRNA in clinical trials has been attributed to a general lack of understanding of host cellular context related to saRNA expression within human immune cells^7,11–13^. *In vitro* and *in vivo* studies have documented a strong interferon response to saRNA^15,16^, and while there is evidence that these same interactions exist in humans, they are currently poorly understood^3^. To improve the potential of saRNA as a therapeutic modality, we aimed to provide a better understanding of the cellular context controlling the regulation of saRNA expression in human cells. To this end, we conducted a CRISPR-KO screen that identified mitochondrial function and ferroptosis as cellular processes that may be essential in and unique to successful host translation and expression of saRNA. Through pharmacological intervention and single gene knockouts, we have demonstrated that this connection to mitochondrial function significantly impacts the expression of saRNA *in vitro*, with mild to no effect on modRNA. While originally screened in immune incompetent 293T cells, we have also shown that our results can be applied to more physiologically relevant cell types such as Jurkat T cells and primary normal human dendritic cells. Furthermore, we have shown that modulation of ACSL4 expression, one of our top hits and a necessary factor of ferroptosis^26^ and single-stranded RNA virus infection^28^, can significantly impact saRNA expression in multiple cell types, but has little to no effect on modRNA expression. To our knowledge, we are the first to demonstrate this integral connection between saRNA expression, mitochondrial function, and ACSL4.

As a vaccine modality, saRNA has been studied in various preclinical models against a wide variety of infectious diseases, most commonly influenza^1,10^. Most promisingly, saRNA has outperformed DNA- and protein-based vaccines in studies of HIV, a notoriously difficult virus to combat^6,8,9^, highlighting the potential of saRNA in vaccines and therapeutics. With minor exception^14^, efforts to realize saRNA as a therapeutic modality have hit many roadblocks with underwhelming results in phase I and II clinical trials of multiple disease types^1,7,11–13^. Although, like modRNA, saRNA is generally well-tolerated with an acceptable safety profile, most saRNA vaccines have been unable to induce an improved immunogenic response or sufficient seroconversion over traditional or modRNA vaccines to date^7,11–13,37^. Most trials reached similar conclusions: that saRNA requires modification “to realize its potential as an effective vaccine”^37^. These setbacks underscore the importance of understanding the host cellular context of saRNA expression and emphasize the significance of our findings in this current study.

To date, studies of saRNA have focused on its interactions with immune factors (e.g. interferons, RIG-I)^3,15,16^. While these factors are undoubtedly essential considerations for adapting saRNA for clinical use, our study highlights that immune machineries are not the only mechanisms by which saRNA is governed and underscores the need to expand upon non-immune regulating factors of expression. Although ferroptosis and mitochondria are gaining interest in immunology, especially with regard to antivirals and cancer vaccines^29,32,34,38–47^, we have documented a need for the expansion on this space with regards to RNA and vaccine modalities. Moreover, because the activity of ferroptotic pathways vary depending on the cell type, especially within immune sub lineages^32^, the successful adaptation of saRNA for therapeutic use requires refinement in order to overcome the limited success of clinical trials to date^7,11–13^.

This work highlights the potential to enhance saRNA efficacy for clinical applications. This could be achieved by modulating ACSL4 in saRNA vaccines, as we have shown that activating ACSL4 by multiple methods consistently improves saRNA expression across multiple cell types without detectable cytotoxicity. Co-delivery of a therapeutic saRNA with ACSL4 modRNA, as we demonstrate in this work, is a viable strategy for adjuvant therapy; one possible continuation of this work is to investigate the duration of the effect of ACSL4 modulation on saRNA expression. Another strategy to increase the efficacy of saRNA could be through nucleotide modification.

Previously, it was widely reported that nucleotide modifications impaired downstream efficacy of saRNA^16,48–53^, therefore all saRNA formulated for this work contained unmodified nucleotides. However, recent evidence suggests alternative nucleotide modifications could improve saRNA efficacy^51^. Incorporating modified nucleotides into saRNA, coupled with ACSL4 activation, could further enhance the efficacy and therefore impact of saRNAs in vaccines and therapeutics. Interestingly, reactivation of ferroptosis mechanisms within tumor cells is gaining interest for antitumor treatments and cancer vaccines^43,46,47^, highlighting another mechanism that may contribute to improving saRNA expression. However, this would require an extremely nuanced approach in order to balance the fine edge between activating the machineries required for saRNA expression and killing target host cells. In summary, our results provide a deeper understanding of the cellular environment governing saRNA expression and regulation, revealing links to key non-immune biological processes, and lay the groundwork for future vaccine and therapeutic applications.

## MATERIALS AND METHODS

### In vitro transcription of RNA

All modRNAs and saRNAs designed and produced in-house were synthesized using in vitro transcription (IVT) protocols described previously^54–56^. Briefly: saGFP and modGFP template plasmid designs were based on standard eGFP and HiBit^20,21^, using vector backbone, UTRs, and polyA length as previously described for Pfizer modRNA design^56–59^. modACSL4 template design was based on published sequences^60^ with Pfizer modRNA backbone, UTRs, and polyA as previously described^56–59^. saRNA replicon was designed as previously described and published^61^. Plasmids were produced by GenScript; NEB® Stable Competent E. coli cells were used for plasmid transformation. Purification and subsequent DNA linearization with BspQI enzyme was performed in-house using manufacturer’s protocols (Qiagen, New England Biolabs respectively). IVT was performed using T7 polymerase and capped co-transcriptionally using CleanCap 5’ cap structures (TriLink). modRNA was generated with N1-Methylpseudouridine-5’-Triphosphate (N1MePsU) and CleanCap AG (both TriLink). saRNA was generated without modified nucleotides and using CleanCap AU (TriLink) based on previous reports that modified nucleotides impair downstream efficacy of saRNA^16,48–53^. saRNA IVT reactions were performed at lower temperatures as previously described in order to improve yields^62^. IVT products were purified post-transcription by DNase incubation and LiCl precipitation before being reconstituted in water. For quality control, RNA was analyzed by spectrophotometer and agarose gel. mod-mCherry was produced by and purchased from TriLink.

### Cell Lines

All human cell lines were sourced from vendors following protocols reviewed and approved by the appropriate regulatory and ethics authorities. 293T cells (ATCC) and fibroblasts (HFF-1, ATCC) were maintained in DMEM (Invitrogen) supplemented with 10% fetal bovine serum (FBS) (Invitrogen). Jurkat derived lines (Clone E6-1; ATCC) were maintained in RPMI-1640 (Invitrogen) with 10% FBS. Normal human dendritic cells (NHDCs; Lonza) were maintained as previously described^63^ in RPMI-1640 with 10% FBS, 1% non-essential amino acids, and 1% GlutaMax (all Invitrogen). All cells were maintained in sterile, humidified incubation at 37°C with 5% CO2.

### Generation of Stable Cas9-expressing Cell Lines

293T-Cas9 cells were generated as previously described^19^. Briefly, 293T cells were transduced with pLenti7-EF1a-Cas9 via lentivirus supplemented with 10 μg/mL polybrene. pLenti7-EF1a-Cas9 lentivirus was designed, produced, and kindly shared with us by our colleagues^19^. 293T-Cas9 cells were selected with 500 µg/mL hygromycin B (Invitrogen) and expanded to create frozen stocks. Cas9 expression and editing efficiency was confirmed by next generation sequencing after lentivirus transduction of a standard housekeeping guide RNA and 1 µg/mL puromycin selection. 293T-dCas9-mCherry cells were generated with the same lentivirus transduction method; dCas9-mCherry plasmid was designed as previously described^36^; lentivirus was produced by our colleagues and kindly shared with us. 293T-dCas9 cells were selected by fluorescence-activated cell sorting (FACS, see below).

### In vitro expression (IVE) assay

293T cells, fibroblasts, and NHDCs were transfected using Lipofectamine MessengerMax (Invitrogen) according to the manufacturer’s protocol. Briefly, cells were seeded 1-2 days prior to transfection with a targeted confluency of 70-80% at transfection. Lipofectamine MessengerMax was incubated with OptiMEM (Invitrogen) at RT prior to addition of RNA (concentrations noted in relevant figures) and further incubation at RT. In the case of co-transfection, RNAs were mixed prior to incubation with MessengerMax. RNA/MessengerMax mixes were added directly to wells and incubated for the time indicated in the relevant figure legend. RNA and MessengerMax concentrations were scaled appropriately based on cell number and culture volume for different culture sizes. Jurkat cells were transfected using Lonza 4D-Nucleofector® in 96-well format according to the manufacturer’s protocol. Briefly, the appropriate number of Jurkat cells were collected, spun down, and resuspended in Nucleofection Buffer SE (Lonza) and appropriate volume of RNA (concentrations specified in relevant figure legends). In the case of co-transfection, RNAs were mixed prior to addition to cells. After nucleofection with Lonza proprietary nucleofection program CL-120, cells were resuspended at relevant density in pre-warmed media and incubated for the time indicated in the relevant figure legend.

IVE results were read by either flow cytometry (GFP) or HiBit luminescence assay (HiBit tag^20,21^). In the case of CRISPR-KO, CRISPRa, and overexpression assay, appropriate number of cells were collected for gene expression assay prior to IVE. In the case of co-transfection of modACSL4, at the time of reporter readout an appropriate number of cells were simultaneously collected for gene expression assay by qPCR (see below). Flow cytometry description is below. HiBit luminescence assay was performed using Nano-Glo® HiBiT Lytic Detection System (Promega) according to manufacturer’s protocol. Briefly, cell media was aspirated from wells before addition of Nano-Glo® HiBiT Lytic buffer supplemented with Nano-Glo® HiBiT Lytic substrate and LgBiT protein. After cells had lifted, cell/buffer mixture was transferred to solid white polystyrene plates (Corning) before reading on EnVision Plate Reader (Perkin Elmer).

### Flow Cytometry

Flow cytometry was performed using standard laboratory protocols. Briefly, cells were lifted as normal (or, in the case of suspension cells, collected and spun down) prior to washing with Dulbecco’s phosphate-buffered saline (DPBS) without Calcium and Magnesium (Invitrogen). Cells were stained with a viability dye of the appropriate fluorophore (BioLegend Zombie Violet™ dye or BD Biosciences Horizon™ Fixable Viability Stain 780) in order not to clash with RNA fluorophores. Cells were fixed with IC Fixation Buffer (Invitrogen) prior to reading on the LSRFortessa™ Cell Analyzer (BD Biosciences); readout was analyzed by FlowJo (BD Biosciences). FACS was performed using standard laboratory protocols. As above, cells were lifted as normal prior to washing with DPBS. Cells were re-suspended in DPBS at appropriate density and maintained on ice before sorting with SH800S Cell Sorter (Sony) and its corresponding software. Cells were maintained in appropriate media on ice prior to downstream processing. FACS readout was also analyzed by FlowJo.

### CRISPR-Cas9 Screening

CRISPR screen was performed as previously described^19^. Briefly, gRNA library was introduced to 293T-Cas9 cells in three separate pools via lentiviral transduction. gRNA library design and lentiviral production was performed as previously described^19^ and lentiviral particles were kindly shared with us. For each gRNA pool, 293T-Cas9 cells were transduced at MOI 0.5 supplemented with 10 μg/mL polybrene in the lentivirus media. 24 hours post-transduction, cells were selected with 1 µg/mL puromycin in addition to 250 µg/mL hygromycin B maintenance for two days. After selection, a representative number of cells were collected as a post-selection control and baseline readout of Cas9 activity. For both control and experimental samples, we collected sufficient cells for a goal library coverage of ≥500 cells/gRNA. Selected cells were subcultured for an additional four days before co-transfection with mod-mCherry (TriLink) or saGFP. Each co-transfection for each pool was repeated the following day in a new batch of cells as a replicate and internal control. After overnight IVE, cells were collected for FACS as described above. Prior to sorting, a representative number of cells were collected as a pre-sort control. During sorting, only the top 50% of GFP+/mCherry+ cells were collected as a representative saRNA sample; all mCherry+ single expressors were collected as a representative modRNA sample. As previously mentioned, no singly GFP+ population existed. Sorted cells were pelleted and frozen at −80°C until batch processing. Genomic DNA was extracted from all cell pellets using Gentra Puregene kit (Qiagen) as per manufacturer’s protocol. First round PCR to amplify genome-integrated gRNA sequences was performed using KOD Hot Start Polymerase (Sigma) as per manufacturer’s protocol. PCR product was purified using QIAquick PCR purification kit (Qiagen) as per manufacturer’s protocol before second round PCR for addition of multiplexing barcodes (i5 and i7; Illumina) was performed using Q5® High-Fidelity kit (New England Biolabs) as per manufacturer’s protocol. Final purified PCR product was sequenced on Illumina NextSeq500. gRNA depletion as a readout for gene knockout was analyzed using Model-based Analysis of Genome-wide CRISPR-Cas9 Knockout (MAGeCK)^64^; each sorted library was compared to its corresponding pre-sort control in order to generate hits. False discovery rate was used as a significance cutoff (FDR≤10%). Only significant hits appearing in both replicate batches were considered “true” hits.

### Pharmacological treatment pre-IVE

All pharmacological agents were purchased as lyophilized powder and resuspended in DMSO according to the manufacturer’s recommendations. 1S,3R-RSL3 (RSL3; Tocris Bioscience), rotenone, antimycin A, and ferrostatin-1 (all Sigma) were diluted in appropriate cell culture media to the final concentrations indicated in relevant figures and added directly to cells for indicated time prior to IVE as described in relevant figure legends.

### Single gene CRISPR-KO and CRISPRa in vitro

For single gene CRISPR-KO: 293T-Cas9 cells generated as described above were thawed and maintained in complete DMEM medium supplemented with 250 µg/mL hygromycin B. Various GOIs were selected based on screening results and pre-designed gRNAs were purchased from IDT and Synthego. 293T-Cas9 cells were transfected with gRNAs using RNAiMAX Lipofectamine (Invitrogen) per manufacturer’s protocol. Briefly, RNAiMAX lipofectamine diluted in OptiMEM was mixed with gRNAs for a final concentration of 25 nM RNA and incubated at RT prior to addition directly to cells at half confluency in a 12 well plate. Cells were incubated for 48-72 hours before being lifted; appropriate numbers of cells were collected for IVE, while remaining were collected for gene knockdown assay. For assaying gene knockdown, cells were pelleted and gDNA was extracted as described above. gDNA was amplified by PCR as described above. Unpurified PCR products were submitted to Azenta for sequencing and analyzed using ICE CRISPR Analysis Tool (Synthego)^65^.

For CRISPRa: 293T-dCas9-mCherry cells generated as described above were used. gRNAs for ACSL4 were designed using CRISPick^66,67^; selected designs were submitted to IDT for production. As above, gRNAs were transfected by RNAiMAX lipofectamine. Cells were incubated for 48 hours before an appropriate number of cells were collected for IVE as described above, and the remaining were collected for gene expression assay by qPCR (see below).

### Single gene overexpression in vitro

Pre-designed pLenti-C-Myc-DDK-P2A-Puro lentiviral particles targeting ACSL4, along with its corresponding control lentivirus particles, were purchased from OriGene. Both were introduced to 293T cells via lentivirus transduction as described above, with a target MOI of 0.3 and 1 µg/mL puromycin selection for 72 hours. After selection, cells were lifted and appropriate number of cells were taken for IVE; remaining cells were frozen as stocks or collected for gene expression assay by qPCR (see below).

### Quantitative Real Time Polymerase Chain Reaction

Total RNA was extracted from cell pellets using appropriate Qiagen RNeasy Plus kit according to the manufacturer’s protocol. cDNA conversion was performed using High Capacity cDNA Reverse Transcription Kit (Applied Biosystems) according to the manufacturer’s protocol. Pre-designed probes for qPCR were purchased from IDT. qPCR was performed using TaqMan™ Fast Advanced kit (Applied Biosystems) and run on QuantStudio 7 Flex (Applied Biosystems) according to the manufacturer’s protocols; all reactions were run in triplicate. Ct values, fold change (2^−ΔΔCt^), and statistical analysis were analyzed in R using package *pcr*^68–70^.

### Lipid Peroxidation Detection

BODIPY Lipid Peroxidation Sensor (Invitrogen) was used as per manufacturer’s protocol. Briefly, BODIPY 665/676 dye (to avoid conflict with GFP) was added directly to wells post-IVE and incubated for 30 minutes in the incubator before collection and flow cytometry prep as normal.

### Quantification and Statistical Analyses

All statistical analyses were performed using GraphPad Prism or R. Statistical tests used were one- or two-way ANOVA with Dunnett’s multiple comparisons test and two-sample *t*-test as appropriate. For the CRISPR-KO screen, FDR<0.1 was used as significance cutoff. For all other data, significance cutoff is p<0.05. All data are reported as mean ± SEM unless otherwise stated.

## Author Contributions

LFT wrote the manuscript and designed this study with supervision from CW and AP. CW originally conceptualized this study with AS, under the guidance of JC-R and LL. LFT performed all experiments with supervision from CW, AP, and SK. QX provided guide RNA library, guidance for CRISPR screen, and performed MAGeCK analysis. AP assisted with manuscript development. SK provided feedback and revisions on the manuscript. CW, AS, QX, JC-R, LL, and AP provided project supervision throughout this study.

## Acknowledgements

We thank the Pfizer Research and Development Postdoctoral Program for their support. We thank former Pfizer colleagues in Emerging Science and Innovation, specifically Chunying Zhao, Meagan Didyk, Deyan Tong, Lisa Kim, and Jason D. Arroyo, for their help with generating reagents, training, transfer of protocols, and assistance with library sequencing. We also thank the Kendall Square Flow Cytometry Shared Resource and its staff Margery Pelletier and Vaughn Youngblood for use of their equipment. We thank Pfizer BioMedicine Design colleague Jonathan McDaniel for training and assistance in library sequencing, and we thank the Kendall Square Next Generation Sequencing Shared Resource for use of their equipment. We would also like to thank past and present Pfizer colleagues Austin M. Krohn, Bianca M. Lauro, and Praveen K. Veerasubramanian for their assistance in generating reagents and providing stimulating discussion on certain aspects of this study.

