## Supplemental Figures for "Self-amplifying mRNA expression is governed by mitochondrial machinery and ACSL4"

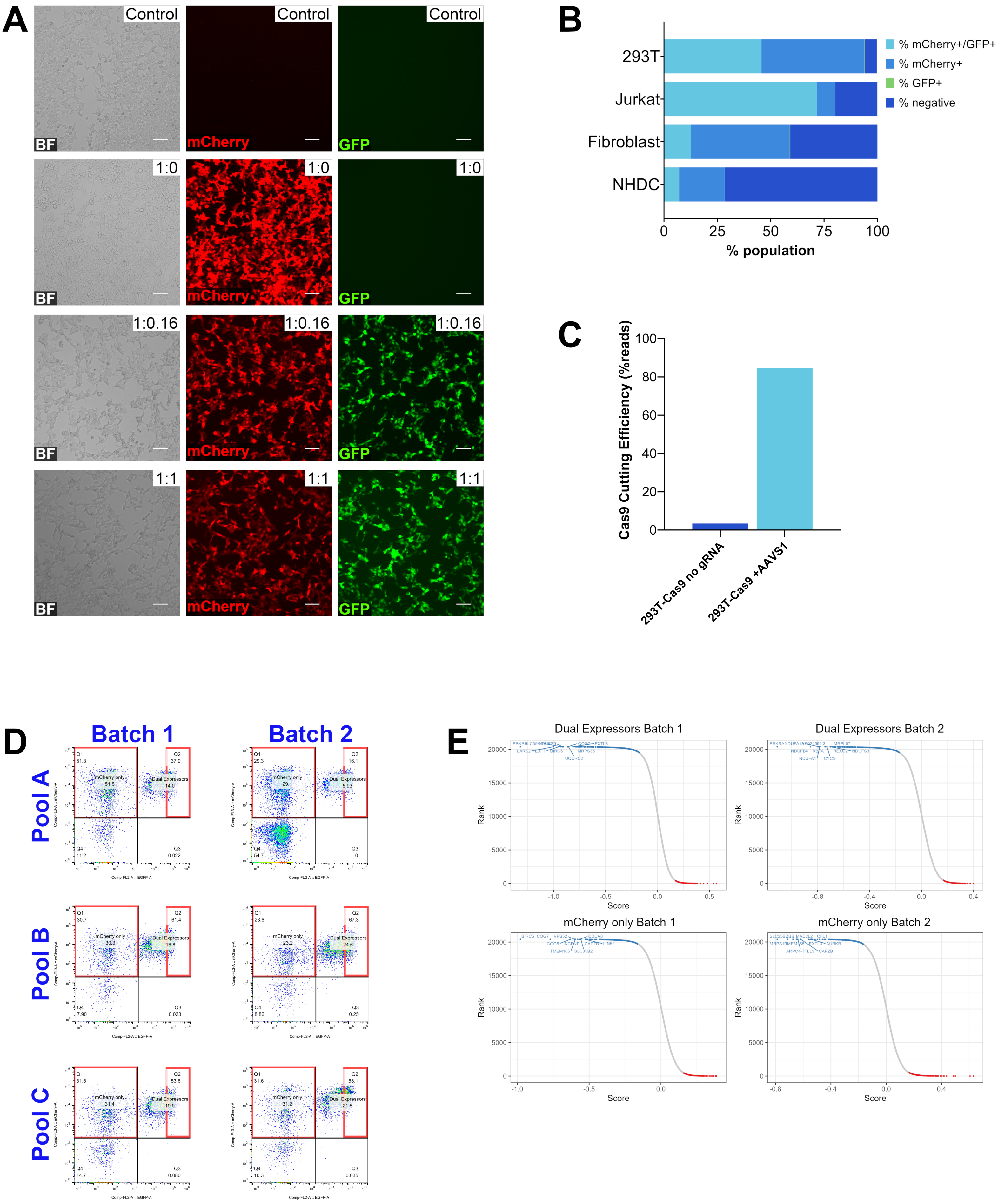
A
Control
1:0
1:0.16
1:1
BF
mCherry
GFP
B
293T
Jurkat
Fibroblast
NHDC
% mCherry+/GFP+
% mCherry+
% GFP+
% negative
0
25
50
75
100
% population
C
Cas9 Cutting Efficiency (%reads)
100
80
60
40
20
0
293T-Cas9 no gRNA
293T-Cas9 +AAVS1
D
Batch 1
Batch 2
Pool A
Pool B
Pool C
mCherry only
Dual Expressors
E
Dual Expressors Batch 1
Dual Expressors Batch 2
mCherry only Batch 1
mCherry only Batch 2
Rank
Score

**SUPPLEMENTAL FIGURE 1: Generating screen challenge parameters, 293T-Cas9 cells, and cell libraries for the CRISPR-KO screen.** A) Representative brightfield, mCherry, and GFP images of 293T cells co-transfected overnight with saGFP and mod-mCherry at the indicated concentration ratio. mod-mCherry concentration was kept constant at 50 ng/well in a 96 well plate while saGFP varied from 4 to 50 ng/well. Images were taken at 10X magnification on the EVOS M5000 (Invitrogen). Scale bars represent 50 µM. B) Overnight co-transfection of saGFP and mod-mCherry in 293T cells, Jurkat cells, fibroblasts, and primary normal human dendritic cells (NHDCs) showing the proportion of live cells that are dual expressing, single expressing, or negative. For 293T and fibroblasts, each well (in 96 well format) received 50 ng/well mod-mCherry and 25 ng/well saGFP. For NHDCs, each well (in 96 well format) received 100 ng/well mod-mCherry and 50 ng/well saGFP. Jurkat cells were nucleofected using Lonza 4D-Nucleofector™ 96-well Unit with a cell density of 5e5 cells per well; each well received 1000 ng mod-mCherry and 500 ng saGFP. Cells were analyzed by flow cytometry; debris, dead cells, and doublets were gated out by FlowJo. C) Cas9 editing efficiency of 293T-Cas9 stable cell lines as shown by percentage of modified sequenced transcripts after introduction of an AAVS1-targeting control gRNA. 293T-Cas9 control cells (left) that received no gRNA compared to edited cells (right). D) Representative flow cytometry scatter plots to show populations sorted and collected for screen libraries (boxed in red) for each Batch and Pool. E) Rank plots for each sample displaying beta score deviation calculated by MAGeCK. Scores help select for top hits; as we focused on negative hits, we focused on genes in the top “negative” scores, highlighted in blue.

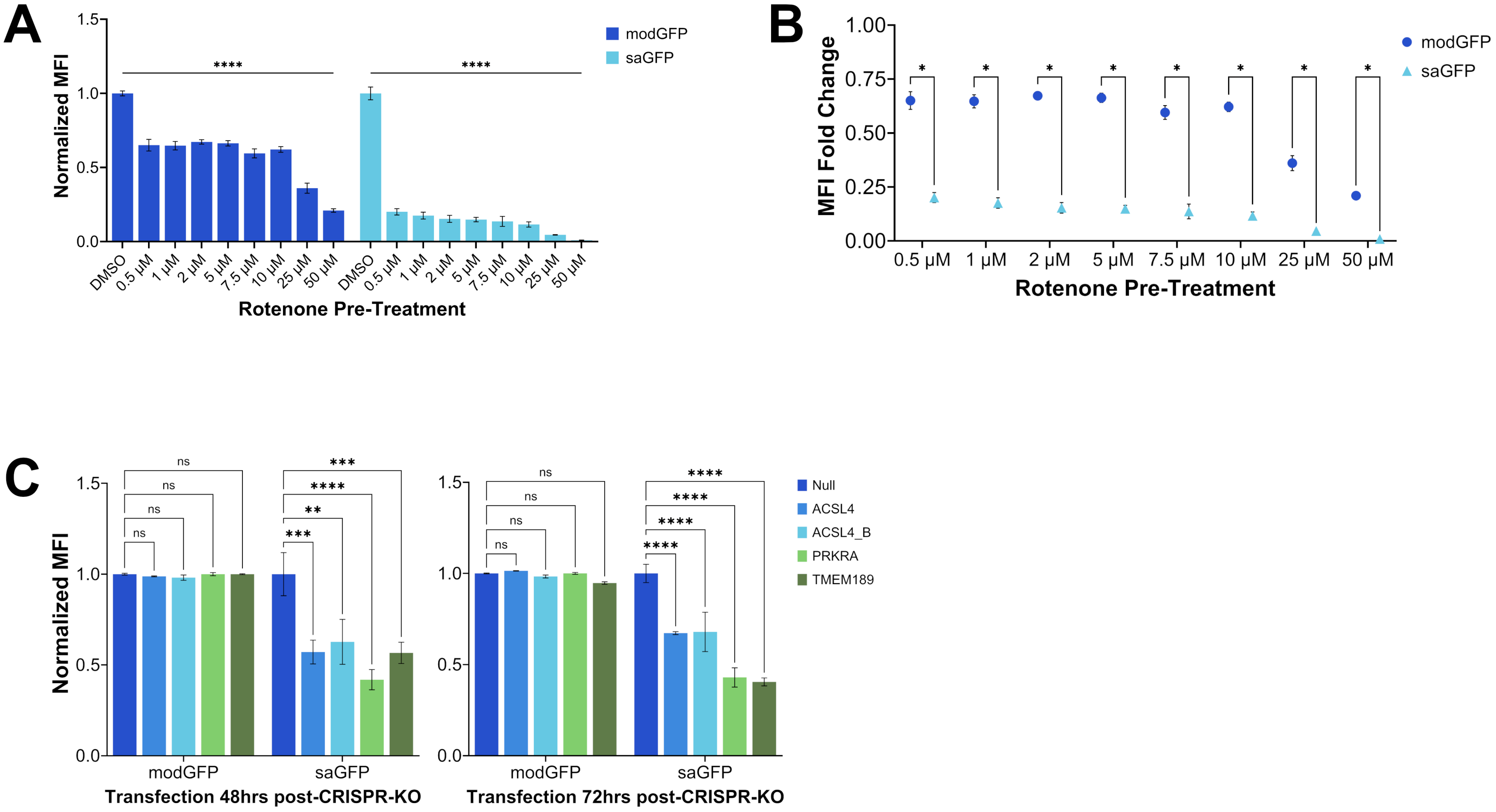

A
Normalized MFI
Rotenone Pre-Treatment
modGFP
saGFP
****
DMSO
0.5 μM
1 μM
2 μM
5 μM
7.5 μM
10 μM
25 μM
50 μM
B
MFI Fold Change
Rotenone Pre-Treatment
modGFP
saGFP
C
Normalized MFI
Null
ACSL4
ACSL4_B
PRKRA
TMEM189
ns
***
****
**
Transfection 48hrs post-CRISPR-KO
Transfection 72hrs post-CRISPR-KO

**SUPPLEMENTAL FIGURE 2: Rotenone treatment and single gene KO show biological processes and genes identified by CRISPR-KO screen are unique to saRNA expression.** A) 293T cells were pre-treated with rotenone overnight at the concentrations indicated before transfection with modGFP or saGFP. Each well received a standard dose of RNA (120 ng/well in 96 well format). MFI was read by HiBit assay after four hours and normalized to the negative control of each modality (untreated/DMSO). n=3 biological replicates. B) Comparison of fold change in MFI for each concentration of rotenone pre-treatment. saRNA MFI is significantly more impacted at all concentrations. C) 293T-Cas9 cells transfected with four guide RNAs targeting hits pulled from various pools including: PRKRA, TMEM189, and two unique ACSL4 guides. KO cells were subcultured for 48 (left) or 72 (right) hours prior to transfection with modGFP or saGFP. Each well received a standard dose of RNA (120 ng/well in 96 well format). MFI was read by HiBit assay after four hours and normalized to the negative control of each modality (gRNA null/normal 293T-Cas9 cells). n=3 biological replicates.

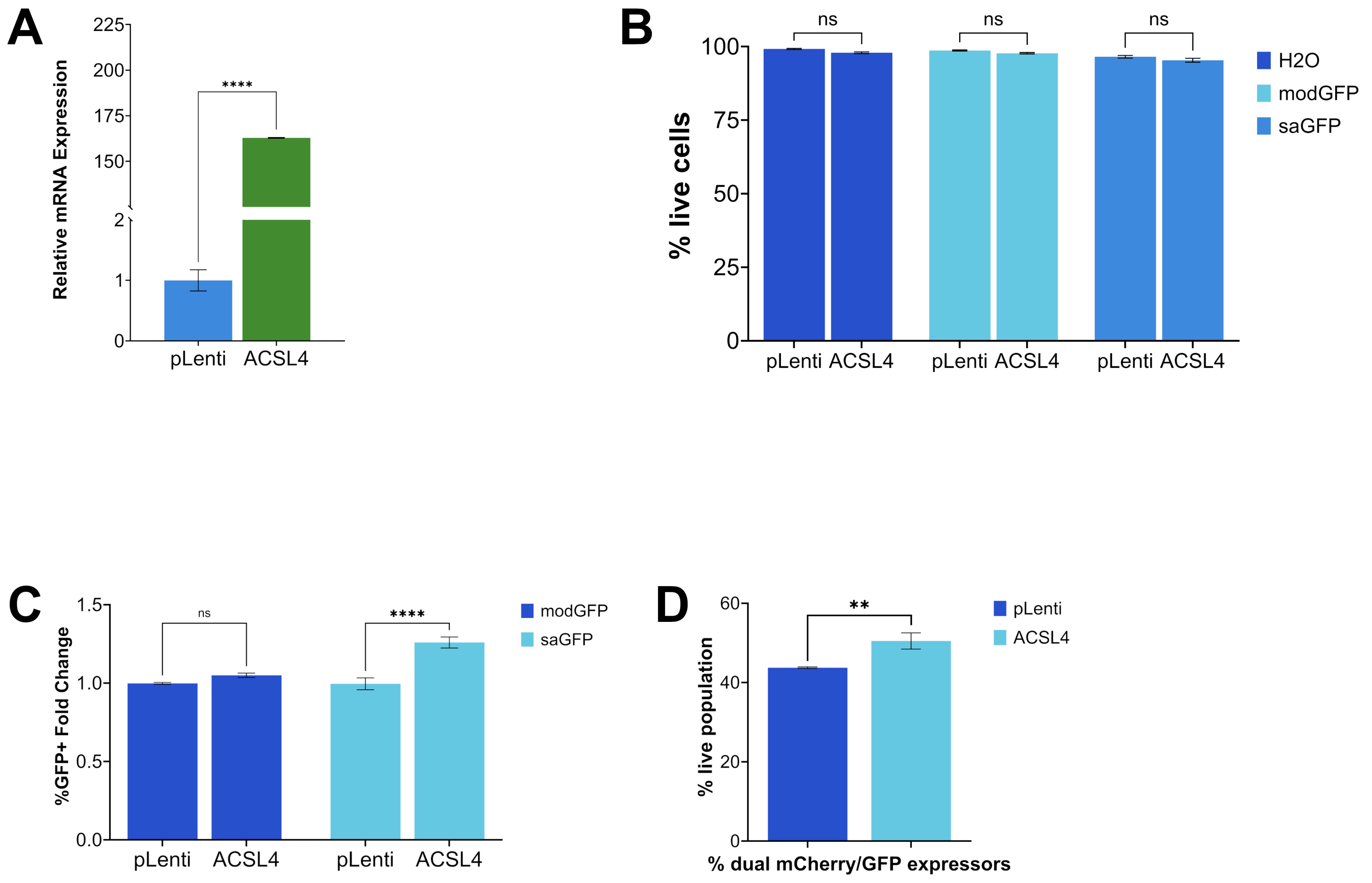
A
Relative mRNA Expression
225
200
175
150
2
1
0
****
pLenti
ACSL4
B
% live cells
100
75
50
25
0
ns
ns
ns
pLenti ACSL4
pLenti ACSL4
pLenti ACSL4
H2O
modGFP
saGFP
C
%GFP+ Fold Change
1.5
1.0
0.5
0.0
ns
****
pLenti
ACSL4
pLenti
ACSL4
modGFP
saGFP
D
% live population
60
40
20
0
**
% dual mCherry/GFP expressors
pLenti
ACSL4

**SUPPLEMENTAL FIGURE 3: ACSL4 overexpression improves saRNA translation without compromising cell viability.** A) ACSL4 expression as read by qPCR shows successful upregulation in 293T cells by overexpression plasmid. Gene expression assay was conducted concurrently with RNA IVE. $2^{-\Delta\Delta CT}$ was calculated using relevant empty Lenti vector (pLenti) cells (lentivirus particles delivered to 293T-Cas9 cells contained no overexpression plasmid) as the control. B) ACSL4 overexpression does not impact viability of cells as seen through proportion of live cells post-transfection. Each well in 96 well format received 100 ng/well RNA, with $H_2O$ as a negative control. Viability was detected by live/dead dye reading after gating out debris and doublets in FlowJo. Each ACSL4 overexpression sample was compared to the pLenti sample that received the same RNA modality. n=3 biological replicates. C) ACSL4 overexpression increases the proportion of GFP+ cells after overnight transfection with modGFP or saGFP. Each well in 96 well format received 100 ng/well RNA, with $H_2O$ as a negative control. Cells were analyzed by flow cytometry; debris, dead cells, doublets, and GFP− cells were gated out by FlowJo to observe GFP+ populations only. Each ACSL4 overexpression sample was compared to the pLenti sample that received the same RNA modality. n=3 biological replicates. D) ACSL4 overexpression increases the proportion of dual expressors after overnight co-transfection with mod-mCherry and saGFP. Each well (in 96 well format) received 50 ng/well mod-mCherry and 25 ng/well saGFP. Cells were analyzed by flow cytometry; debris, dead cells, and doublets were gated out by FlowJo. ACSL4 overexpression samples were compared to a corresponding pLenti sample. n=3 biological replicates.

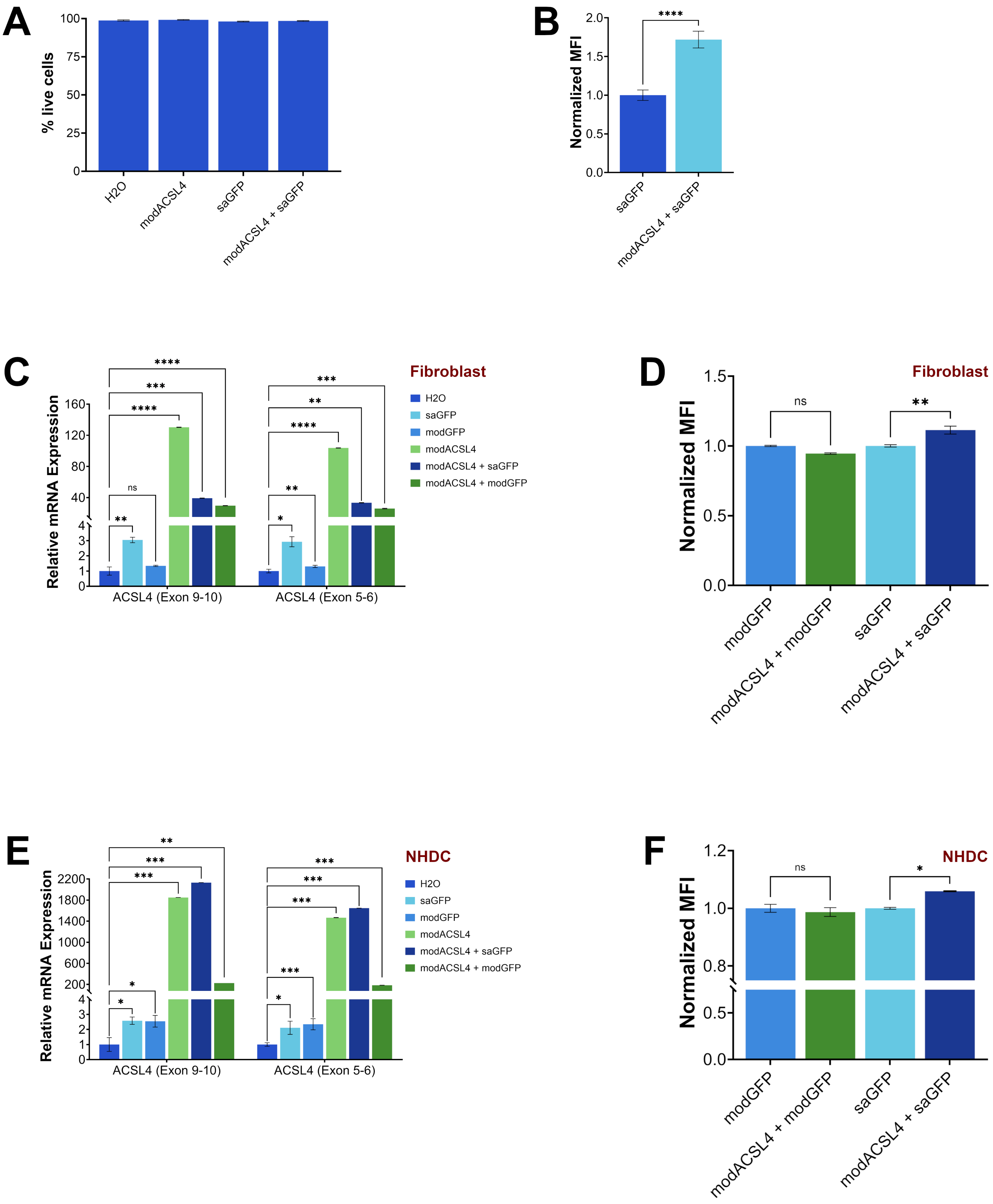
A
% live cells
100
75
50
25
0
H2O
modACSL4
saGFP
modACSL4 + saGFP
B
Normalized MFI
2.0
1.5
1.0
0.5
0.0
****
saGFP
modACSL4 + saGFP
C
Relative mRNA Expression
160
120
80
40
4
3
2
1
0
****
***
****
ns
**
***
**
****
**
*
ACSL4 (Exon 9-10)
ACSL4 (Exon 5-6)
Fibroblast
H2O
saGFP
modGFP
modACSL4
modACSL4 + saGFP
modACSL4 + modGFP
D
Fibroblast
Normalized MFI
1.5
1.0
0.5
0.0
ns
**
modGFP
modACSL4 + modGFP
saGFP
modACSL4 + saGFP
E
Relative mRNA Expression
2200
1800
1400
1000
600
200
4
3
2
1
0
**
***
***
*
*
***
***
***
***
*
ACSL4 (Exon 9-10)
ACSL4 (Exon 5-6)
NHDC
H2O
saGFP
modGFP
modACSL4
modACSL4 + saGFP
modACSL4 + modGFP
F
NHDC
Normalized MFI
1.2
1.0
0.8
0.5
0.0
ns
*
modGFP
modACSL4 + modGFP
saGFP
modACSL4 + saGFP

**SUPPLEMENTAL FIGURE 4: Activating ACSL4 by co-delivery of ACSL4-targeting modRNA and saRNA improves saRNA expression in multiple cell types.** A) 293T cells transfected overnight in a 96 well format with modACSL4, saGFP, or co-delivered both, do not exhibit differences in proportion of live cells as read by flow cytometry. Each single modality transfection received 100 ng/well RNA. Co-transfection with saRNA received 50 ng modRNA/well combined with 25 ng saRNA/well. Viability was detected by live/dead dye reading after gating out debris and doublets on FlowJo. n=3 biological replicates. B) 293T cells co-transfected with modACSL4 and saGFP show increased saRNA expression compared to saGFP transfected alone. Cells were transfected as described in S4A and analyzed by flow cytometry; debris, dead cells, doublets, and GFP− cells were gated out by FlowJo to observe MFI in GFP+ cells only. MFI of each co-transfection was normalized to its single modality counterpart. n=3 biological replicates. C) Transfection with modACSL4 successfully increases ACSL4 expression in fibroblast HFF-1 cells as read by qPCR. Fibroblasts were transfected with either modGFP, saGFP, modASCL4, or co-delivered modACSL4 with a GFP carrying RNA overnight in a 96 well format. Each single modality transfection received 100 ng/well RNA. Co-transfection with saRNA received 50 ng modRNA/well combined with 25 ng saRNA/well. Co-transfection of two modRNAs received 50 ng of each modRNA per well. All cells that received modACSL4 (either alone or co-delivered) exhibited significantly higher ACSL4 expression with both unique probes. $2^{-\Delta\Delta CT}$ was calculated using non-transfected 293T cells ($H_2O$ negative control) as the control. D) MFI of fibroblasts with co-delivered modACSL4 and GFP RNA show increased saRNA expression with no effect on modRNA expression. Fibroblasts were transfected with either modGFP, saGFP, modASCL4, or co-delivered modACSL4 with a GFP carrying RNA overnight in a 384 well format. Each single modality transfection received 50 ng/well RNA. Co-transfection with saRNA received 25 ng modRNA/well combined with 12.5 ng saRNA/well. Co-transfection of two modRNAs received 25 ng of each modRNA per well. MFI of each co-transfection was normalized to its single modality counterpart. n=4 biological replicates. E) Transfection with modACSL4 successfully increases ACSL4 expression in normal human dendritic cells (NHDCs) as read by qPCR. NHDCs were transfected with either modGFP, saGFP, modASCL4, or co-delivered modACSL4 with a GFP carrying RNA overnight in a 96 well format. Each single modality transfection received 200 ng/well RNA. Co-transfection with saRNA received 100 ng modRNA/well combined with 50 ng saRNA/well. Co-transfection of two modRNAs received 100 ng of each modRNA per well. All cells that received modACSL4 (either alone or co-delivered) exhibited significantly higher ACSL4 expression with both unique probes. $2^{-\Delta\Delta CT}$ was calculated using non-transfected 293T cells ($H_2O$ negative control) as the control. F) MFI of NHDCs with co-delivered modACSL4 and GFP RNA show increased saRNA expression with no effect on modRNA expression. NHDCs were transfected with either modGFP, saGFP, modASCL4, or co-delivered modACSL4 with a GFP carrying RNA overnight in a 384 well format. Each single modality transfection received 100 ng/well RNA. Co-transfection with saRNA received 50 ng modRNA/well combined with 25 ng saRNA/well. Co-transfection of two modRNAs received 50 ng of each modRNA per well. MFI of each co-transfection was normalized to its single modality counterpart. n=2 biological replicates.
